# Variability and Landmark Quality in Anatomical Co-Registration in Neuronavigation for Brain Stimulation

**DOI:** 10.64898/2026.10.24.761558

**Authors:** Zhino Ebrahimi, Maryam Farahmandrad, Behnam Seyedi, Ziye Xie, J. Matias Di Martino, Angel V. Peterchev, Thomas Kammer, Stephan M Goetz

## Abstract

Accurate neuronavigation depends not only on the technical performance of the tracking system but also on the reproducibility with which anatomical landmarks are physically localized. We quantified operator- and landmark-specific variability in repeated facial and cranial landmark localization in 19,630 measurements from 16 participants, four operators, and 13 landmarks under marked and unmarked conditions. Within-operator precision, inter-operator agreement, and marked–unmarked displacement were analyzed separately. Median within-operator precision was 1.60 mm and median inter-operator distance was 2.91 mm, with substantial differences across both operators and landmarks. The left medial canthus showed the greatest within-operator precision (1.43 mm), whereas the pronasale showed the closest inter-operator agreement (2.05 mm). Marking the positions before the procedure did not significantly improve within-operator precision overall but reduced inter-operator distances by 29.2%, which indicates convergence on a shared spatial reference rather than improved intrinsic repeatability. Between-operator differences also contained systematic directional components; signed offsets approached 1.5 mm. Exploratory analyses did not suggest general left–right difference in precision, although the approach side produced systematic spatial shifts for midline landmarks despite comparable repeatability. These findings demonstrate that anatomical landmark localization is both operator- and landmark-specific. Standardized landmark definitions, operator calibration, consistent pointer approach, and the use of multiple well-defined registration features should therefore be considered integral components of neuronavigation quality control.

## 1. Introduction

Focused brain stimulation, such as transcranial magnetic stimulation (TMS) or focused ultrasound stimulation (FUS), relies on accurate targeting and placement of the transducer. Transducer position and orientation shape the delivered field distribution and therefore determine the activated cerebral circuits. Conventional TMS coil positioning and heuristic target localization are acknowledged sources of variability [1, 2, 3, 4]. Differences in stimulation location have also been associated with differences in behavioral and clinical outcomes [5, 6, 7]. This spatial problem has become increasingly important as stimulation protocols move toward more focal, individualized, and network-informed targets [8, 9, 10].

Neuronavigation can reduce some of this variability. It can improve the reproducibility of transducer positioning compared with conventional scalp-based approaches and the maintenance of the position [4, 2, 11]. However, the implied precision of the numbers displayed with micrometer resolution can obscure accuracy limitations. The technology underlying most frameless stereotaxy systems was adapted from coordinate measurement systems [12]. Their technical tracking performance, however, does not by itself characterize the quality of the anatomical co-registration.

Recently, Riis et al. compared individual-anatomy-based with template-based targeting of deep brain structures here in *Brain Stimulation* [13]. Such targeting approaches depend on co-registration and accurate matching of landmarks. Previous studies have likewise analyzed end-to-end neuronavigation accuracy and several of its component error sources [4, 14, 15, 16]. Despite extensive analyses of end-to-end neuronavigation accuracy and its individual error sources, the manual localization of facial and cranial landmarks for co-registration in noninvasive brain stimulation remains comparatively under-characterized. As a critical manual step in co-registration, which maps the participant’s physical head anatomy to image space and the virtual head model, an operator identifies physical anatomical landmarks on the participant and records their coordinates. Prominent accuracy analyses in the brain stimulation neuronavigation literature have estimated or simulated the error associated with this step rather than directly quantifying its operator-dependent reproducibility [17, 14]. Related surgical studies have quantified the scatter of landmark selections and shown that localization variability differs across landmarks and operator experience [18, 19]. However, such repeatability measures do not directly quantify systematic differences between operators in where the same physical landmark is localized. Classical point-based registration-error formulations commonly use simplified isotropic assumptions for fiducial localization error [20], although such uniformity across anatomical landmarks appears unlikely.

Accordingly, the manual landmark-localization step remains insufficiently characterized for brain stimulation, including several open questions: (1) How consistently can operators identify the same anatomical reference points, (2) how strongly does localization depend on the operator, (3) are there systematic spatial biases between operators, and (4) which anatomical landmarks provide the most reproducible reference points and would be preferable for brain stimulation?

We therefore isolated the physical landmark-localization step itself. We asked whether repeated localization depends on the operator and the anatomical landmark, whether within-operator precision and inter-operator agreement capture the same component of spatial variability, and whether visible landmark marking reduces either source of variability. In 19,630 eligible measurements from 16 participants and four operators, we directly quantified operator- and landmark-specific variability in anatomical landmark localization.

## 2. Methods

We studied 16 university students (14 women; *M*_age_ = 25.88 years, *SD* = 3.52, range 22–33), who received € 15 as compensation. The study was approved by the Ethics Committee of RPTU (No. SOWI84). Coordinates were tracked using an NDI Polaris Lyra camera and recorded in Brainsight neuronavigation software (Rogue Res., Montreal Canada) for subsequent analysis. A low-weight tracker with flat disk reflectors (Brainsight) attached to the forehead through electromyography electrodes served as subject reference. A handheld pointer (P-1767) served for digitization. In the right-handed coordinate system, *x* runs from right to left, *y* from anterior to posterior, and *z* from inferior to superior. Four right-handed operators repeatedly localized 13 facial and cranial landmarks under marked and unmarked conditions (see below) where available, yielding 19,630 eligible measurements. Incomplete cells were retained as missing rather than imputed.

Study landmarks comprised nasion, pronasale, subnasale, bilateral medial and lateral canthi, bilateral alar curvature points, bilateral posterior points of the anterior auricular notch (*incisura anterior auris posterior*), and bilateral inferior points of the intertragic notch (*incisura intertragica inferior*) (Fig. 1). Landmark terminology was based on established craniofacial anthropometric nomenclature [21]. For the marked condition, Operator 4 marked all anatomical locations with a skin-friendly pen before measurements, and the same visible marks were subsequently used by all operators. In the unmarked condition, operators localized the anatomical landmarks independently without visible markings. Acquisition order was randomized.

**Figure 1:**
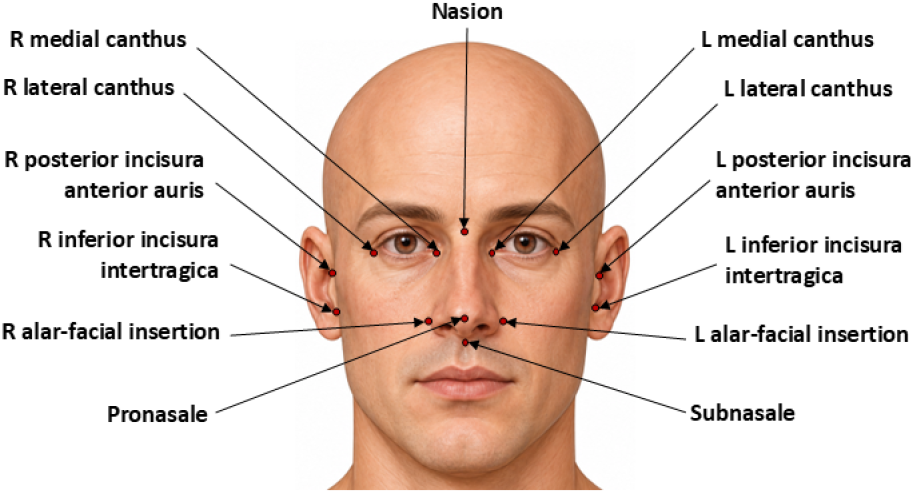
Anatomical locations of the 13 facial and cranial landmarks used in the study. English landmark names are shown in the figure; the corresponding anthropometric terminology is defined in the Methods.

The final analysis comprised 1,586 within-operator precision cells, 2,340 inter-operator distances, and 767 marked–unmarked displacements. Within-operator precision was the median Euclidean distance of repeated measurements from their geometric-median center. Inter-operator agreement was the distance between operator-specific centers, and marked–unmarked displacement was the distance between corresponding condition-specific centers. Log-distance linear mixed-effects models tested operator, marking, operator-pair, landmark, and relevant interactions; estimates were back-transformed to millimeters and planned comparisons were Holm-corrected. For an exploratory assessment of systematic between-operator directional offsets, signed differences between operator-specific landmark centers were averaged across landmarks within participant and examined separately along the three coordinate axes using Wilcoxon signed-rank tests with Holm correction. For the subnasale, marked–unmarked displacement was additionally decomposed into absolute *x, y*, and *z* components, with participant-level aggregation used as a sensitivity analysis. Acquisition order and its interaction with operator were examined on an exploratory basis.

Laterality was examined in two complementary exploratory analyses. First, anatomical laterality was assessed across the five bilateral landmark families by comparing homologous left- and right-sided landmarks; nasion, pronasale, and subnasale were excluded from this comparison because they are anatomically midline. Participant-level left–right differences in within-operator precision and inter-operator distance were assessed using paired Wilcoxon signed-rank tests, with Holm correction for landmark-specific comparisons, and side dependence of signed operator offsets was examined separately by operator pair and coordinate axis. Second, Operator 1 localized the three midline landmarks from both sides of the participant. Therefore, these measurements were additionally classified by approach side. Left- and right-approach measurements were compared for within-approach precision and for signed displacement between approach-specific landmark centers, with Holm correction across the three coordinate axes.

## 3. Results

Within-operator localization was generally precise, but precision varied substantially across operators and landmarks. Median within-operator precision was 1.60 mm (IQR 1.25–2.04; Fig. 2). Operator differences were substantial, *X*^2^ (3) = 356.13, *p* < .001: adjusted precision distance was smallest for Operator 1 (1.31 mm), followed by Operator 2 (1.47 mm), Operator 4 (1.74 mm), and Operator 3 (1.89 mm), with all pairwise comparisons significant after Holm correction (Fig. 2). Precision also differed across landmarks, *X*^2^ (12) = 132.06, *p* < .001. The left medial canthus showed the greatest within-operator precision (1.43 mm), the subnasale the lowest (2.03 mm; Fig. 5). The exploratory operator-by-landmark interaction was significant, *X*^2^ (36) = 186.44, *p* < .001; operator effects were significant within all 13 landmarks after Holm correction. Inter-operator agreement showed a different pattern. Median inter-operator distance was 2.91 mm and differed between operator pairs, *X*^2^ (5) = 41.19, *p* < .001 (Fig. 3), and landmarks, *X*^2^ (12) = 122.58, *p* < .001 (Fig. 5). The pronasale showed the smallest adjusted inter-operator distance (2.05 mm), the right lateral canthus the largest (3.71 mm; Fig. 5). Thus, the landmark with the poorest within-operator precision was not the landmark producing the greatest inter-operator disagreement. Together, these outcomes capture distinct components of spatial reproducibility.

**Figure 2:**
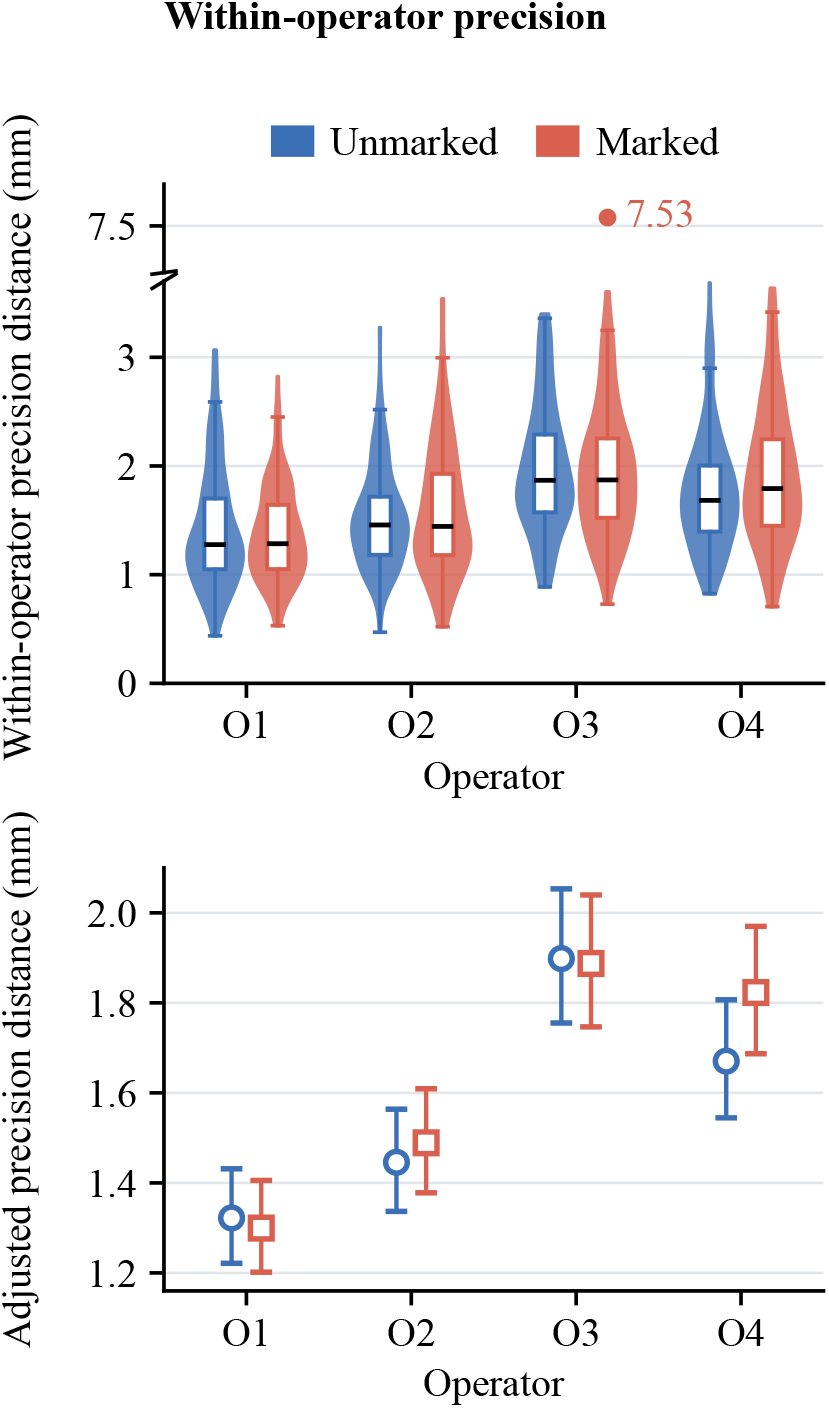
Within-operator localization precision. Top: observed precision-distance distributions for marked and unmarked measurements. Bottom: model-adjusted precision distances by operator and marking condition. Error bars in the model-adjusted panel indicate 95% confidence intervals. Lower distances indicate greater within-operator precision.

**Figure 3:**
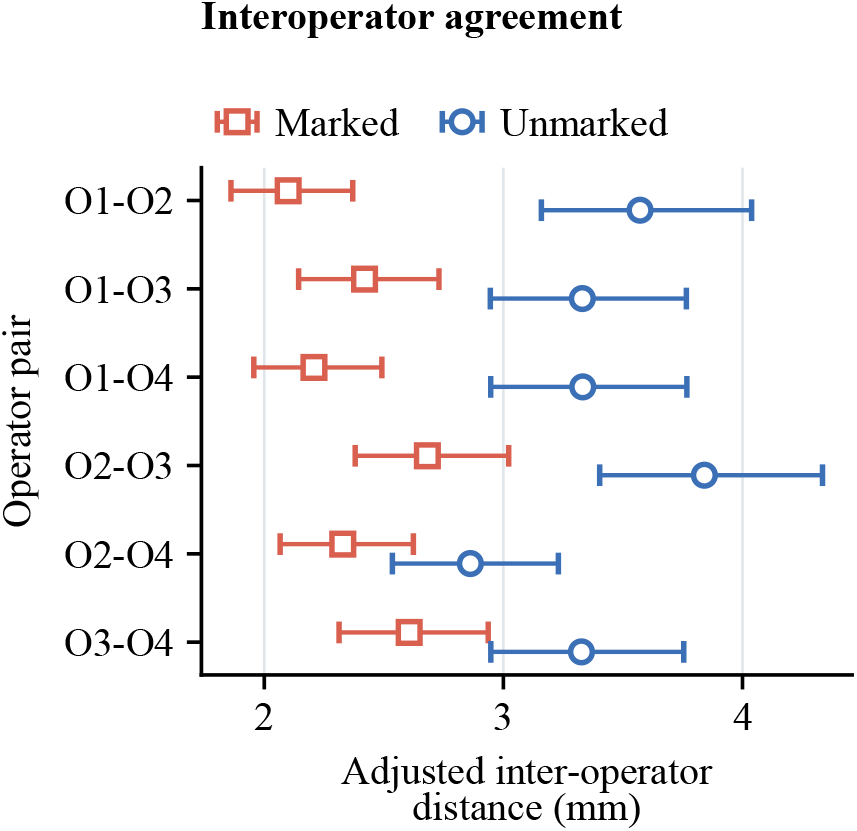
Model-adjusted inter-operator agreement for each operator pair under marked and unmarked conditions. Points represent adjusted inter-operator distances and error bars indicate 95% confidence intervals. Lower distances indicate closer inter-operator agreement.

Inter-operator disagreement was also directional. Exploratory analysis of signed operator-center differences revealed systematic spatial offsets, particularly along the superior–inferior axis. The clearest persistent offset occurred between Operators 3 and 2: Operator 3 localized landmarks on average 1.46 mm inferior to Operator 2 in the unmarked condition (*p*_Holm_ = .001), and the difference remained 1.10 mm under marking (*p*_Holm_ = .003). An unmarked offset of 1.49 mm was also observed between Operators 2 and 4 (*p*_Holm_ = .001), but it decreased to 0.34 mm under marking and was no longer significant. Thus, between-operator disagreement included systematic directional components in addition to overall spatial separation.

Marking did not improve within-operator precision overall, *X*^2^(1) = 2.36, *p* = .125, and the operator-by-marking interaction was not significant, *X*^2^ (3) = 7.28, *p* = .064. The exploratory landmark-by-marking interaction was significant, *X*^2^ (12) = 42.01, *p* < .001. After Holm correction, only the nasion offered a significant landmark-specific benefit: adjusted precision distance was 1.29 mm marked versus 1.64 mm unmarked, a 21.7% reduction. In contrast, inter-operator distances were 29.2% smaller when landmarks were marked, *X*^2^ (1) = 243.64, *p* < .001. The operator-pair-by-marking interaction was also significant, *X*^2^ (5) = 24.13, *p* < .001, and all six operator pairs showed smaller marked distances after Holm correction (Fig. 1C).

Because Operator 4 established the visible marks for all participants, this finding is most consistent with convergence on a common spatial reference rather than improved intrinsic repeatability of individual operators. Marked–unmarked displacement also differed between operators, *X*^2^(3) = 25.42, *p* < .001, and landmarks, *X*^2^ (12) = 46.64, *p* < .001 (Figs. 4 and 5). Adjusted displacement ranged from 2.10 mm for Operator 1 to 2.78 mm for Operator 2. Operator 1 achieved significantly smaller displacement than Operators 2, 3, and 4 after Holm correction, whereas the latter three operators did not differ significantly from one another. Across landmarks, displacement was smallest at the left medial canthus (1.84 mm) and largest at the right posterior point of the anterior auricular notch (3.15 mm). Acquisition order was exploratory: a one-SD later sequence position was associated with approximately 4.5% smaller precision distance (*p* = .014), but the operator-by-order interaction was not significant (*p* = .938) and no operator-specific slope survived Holm correction. We therefore did not interpret this as evidence of learning.

**Figure 4:**
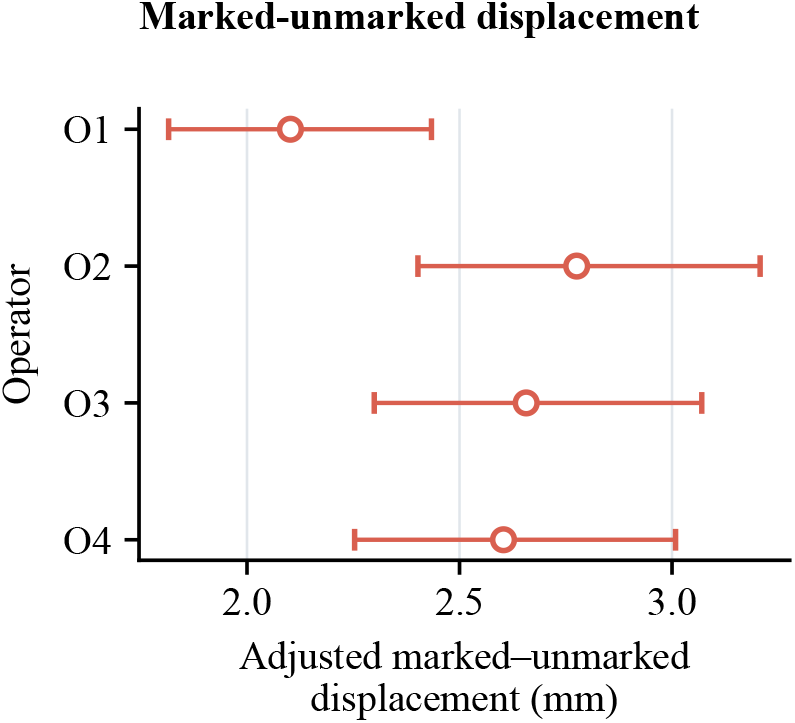
Model-adjusted marked–unmarked displacement by operator. Points represent adjusted displacement estimates and error bars indicate 95% confidence intervals. Lower values indicate a smaller spatial shift between marked and unmarked conditions.

**Figure 5:**
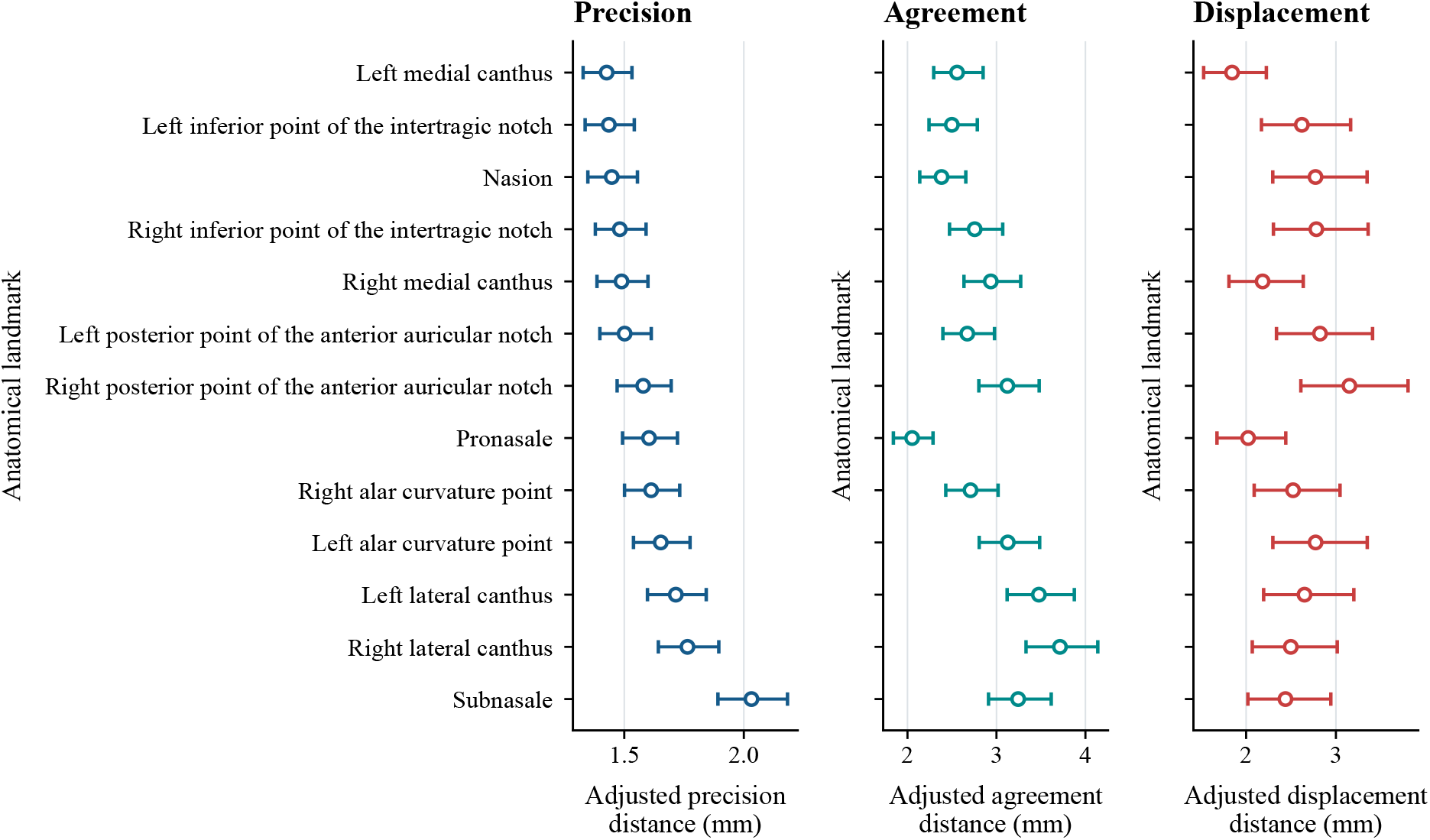
Landmark-specific model-adjusted estimates of within-operator precision, inter-operator agreement, and marked–unmarked displacement. The same landmark order, based on increasing adjusted precision distance, is used across all three outcomes. Points represent adjusted estimates and error bars indicate 95% confidence intervals. Lower precision and agreement distances indicate greater within-operator precision and closer inter-operator agreement, respectively; lower displacement values indicate a smaller spatial shift between marked and unmarked conditions.

Landmark performance was strongly location-specific and, importantly, depended on the aspect of reproducibility considered. The left medial canthus demonstrated the greatest within-operator precision (1.43 mm) and the smallest marked–unmarked displacement (1.84 mm), whereas the pronasale offered the closest inter-operator agreement (2.05 mm). In contrast, the subnasale showed the poorest within-operator precision (2.03 mm), the right lateral canthus the greatest inter-operator disagreement (3.71 mm), and the right posterior point of the anterior auricular notch the largest marked–unmarked displacement (3.15 mm). Thus, there was no single universally *superior* or *bad* landmark; repeatability, agreement, and condition-dependent stability captured complementary properties of landmark performance.

Although no landmark was optimal across every outcome, some locations showed more favorable overall profiles. The medial canthi, particularly the left medial canthus, combined high repeatability with low condition-dependent displacement, whereas the pronasale demonstrated the closest inter-operator agreement. In contrast, the subnasale was comparatively poorly repeatable, the lateral canthi, particularly on the right, showed greater inter-operator disagreement, and posterior points of the anterior auricular notch caused relatively large marked–unmarked displacement.

The subnasale was particularly informative. Its comparatively poor repeatability was accompanied by a directional structure in marked–unmarked displacement: the median absolute *y* component was 1.27 mm, compared with 0.77 mm in *x* and 0.99 mm in *z*. This directionality of the error suggests an influence of the local landmark geometry and/or soft-tissue deformation due to pressure during pointer placement. However, the axis-specific effect was attenuated after aggregation at the participant level; so the mechanical explanation should be considered hypothesis-generating rather than established.

For soft-tissue landmarks such as the subnasale, this pattern suggests that pointer contact, pressure, and approach angle (ideally fully perpendicular for example, which is not possible due to the optics for many targets) should be standardized carefully. In particular, future protocols should minimize and standardize contact pressure and should test whether low-force or contact-controlled localization improves reproducibility at deformable anatomical sites.

The auricular landmarks uncovered another practically relevant pattern. Descriptively, the inferior points of the intertragic notch offered better repeated-localization performance than the corresponding posterior points of the anterior auricular notch, and the right posterior point of the anterior auricular notch showed the largest marked–unmarked displacement of all 13 landmarks. This displacement suggests that proximity to a visually salient anatomical structure does not necessarily translate into superior spatial reproducibility and that the exact definition of the auricular reference point matters.

Exploratory laterality analyses did not reveal a general effect of the body side on within-operator precision. Across the five bilateral landmark pairs, median precision was 1.56 mm on the left and 1.59 mm on the right (geometric left/right ratio = 0.983; *W* = 50, *p* = .375), and no landmark-specific precision contrast survived Holm correction. Overall inter-operator-distance magnitude likewise did not differ significantly between sides (*p* = .159), although signed operator offsets showed side-dependent effects for several operator-pair and coordinate-axis combinations. Inter-operator disagreement at the medial canthus was 0.59 mm smaller on the left (*p*_Holm_ = .011). Thus, the descriptive left–right pattern in precision did not reflect a general anatomical laterality effect.

Operator 1 provided a complementary within-operator test of approach side because nasion, pronasale, and subnasale were localized from both sides of the participant. Within-approach precision was similar for left- and right-sided approaches (median 1.11 vs. 1.16 mm; geometric left/right ratio = 0.977; *p* = .561). Nevertheless, the resulting landmark centers differed systematically: relative to right-sided approaches, left-approach centers were shifted on average 2.11 mm leftward and 1.24 mm superior, with both components surviving Holm correction (*p*_Holm_ < .001); the anterior–posterior component was not significant (*p* = .229). The median three-dimensional separation between approach-specific centers was 2.90 mm.

## 4. Discussion

Together, these findings underline that the reproducibility of landmark-localization cannot be reduced to a single system-level value. Unlike work that estimates registration error indirectly or evaluates downstream coil placement, the present data isolate the manual physical landmark-localization step itself. Anatomical landmark localization was operator-specific and landmark-specific, and repeatability within an operator was distinct from agreement between operators. Moreover, some between-operator differences were systematic in direction rather than merely greater in magnitude, with signed offsets approaching 1.5 mm between particular operator pairs. The persistence of one such offset despite visible marking indicates that a shared reference can reduce overall disagreement without necessarily eliminating operator-specific localization tendencies, whereas the marked attenuation of another offset shows that some biases can be constrained substantially by a common physical reference. A common visible reference reduced inter-operator distances by 29.2% without improving within-operator precision overall, which suggests standardization of the shared spatial reference rather than improved intrinsic repeatability. Training, written landmark definitions, operator calibration, and documentation of marking procedures should therefore be treated as quality-control components of neuronavigated brain stimulation.

The landmark-specific findings also suggest that registration geometry deserves consideration. Because most facial landmarks occupy a comparatively anterior and near-planar configuration, well-defined auricular landmarks may provide useful additional spatial constraint away from the facial *xz* plane. For auricular registration, these findings argue against relying on a single loosely defined auricular reference. When the protocol permits, recording both posterior points of the anterior auricular notch and inferior points of the intertragic notch bilaterally may provide multiple off-plane spatial constraints and help stabilize the anterior–posterior dimension of the registration geometry. Because the posterior points of the anterior auricular notch were in comparison less reproducible, particularly on the right, the contribution of individual auricular landmarks may ultimately need to be optimized rather than treated as equivalent. The present study did not directly compare target registration error across alternative landmark configurations, so the benefit of such multi-point auricular configurations remains to be tested directly.

A complementary approach is surface-based registration, in which multiple scalp points are used to match the participant’s head surface to a head model. Nieminen et al. [14] compared landmark- and surface-based head-to-MRI registration and reported lower registration errors for the surface-based approach. Their implementation sampled points distributed across 21 scalp regions rather than relying on a small set of discrete anatomical landmarks. Surface-based registration may therefore reduce dependence on the localization of individual landmarks when an appropriate head model is available.

The laterality analyses further distinguish repeatability from systematic localization tendency. Anatomical side did not produce a general difference in within-operator precision. More revealingly, Operator 1 reached comparable precision when approaching the same midline landmarks from opposite sides, i.e., the operator could reliably position the pointer tip at the same position for that specific approach side; yet the resulting landmark centers were systematically displaced, i.e., the points for the left approach and the right approach differed. Approach side, viewing geometry, pointer orientation, or operator posture may therefore alter the estimated location of a landmark without measurably reducing within-side repeatability. Because this approach-side comparison was available for only one operator, and all operators were right-handed, the effect cannot be attributed specifically to handedness and should be considered exploratory.

In synopsis, these findings support several practical recommendations for landmark-based neuronavigation. Anatomical landmarks should not be treated as interchangeable: the medial canthi provide comparatively reproducible references, whereas the subnasale suffered from poorer repeatability and therefore requires particular attention to contact pressure and approach geometry. The pronasale offered the closest inter-operator agreement, whereas the right lateral canthus in comparison led to greater inter-operator disagreement. When multiple operators are involved, a shared visible reference can substantially reduce inter-operator disagreement, but should be accompanied by standardized landmark definitions, operator training, and calibration. Approach side and pointer geometry should likewise be standardized or documented, because systematic spatial shifts can occur even when repeatability remains unchanged. Finally, for registration geometry, bilateral posterior points of the anterior auricular notch and inferior points of the intertragic notch would allow stabilizing and reducing the error of the orientation and the scaling of the *y* axis. As all other points are almost fully in the *xz* plane established by the face, multiple auricular points could compensate that imbalance and average out the error of just a single auricular reference. In general, we recommend co-registering with substantially more features than the minimum, which is usually used to date, and the ones tested here provide a catalog as well as corresponding reproducibility features for each. These procedural factors should be considered part of neuronavigation quality control rather than assumed to be negligible relative to the technical accuracy of the tracking system.

## Notes

### Competing Interest Statement

The authors are inventors of brain stimulation technology owned by current and previous employers.

### Summary of Updates

Typographical errors in author list corrected

## References

[1] M. Koehler, T. Kammer, S. M. Goetz, How coil misalignment and mispositioning in transcranial magnetic stimulation affect the stimulation strength at the target, Clinical Neurophysiology 162 (2024) 159–161. doi:10.1016/j.clinph.2024.03.037.

[2] U. Herwig, F. Padberg, J. Unger, M. Spitzer, C. Schönfeldt-Lecuona, Transcranial magnetic stimulation in therapy studies: examination of the reliability of “standard” coil positioning by neuronavigation, Biological Psychiatry 50 (1) (2001) 58–61. doi: 10.1016/S0006-3223(01)01153-2.

[3] A. Mir-Moghtadaei, R. Caballero, P. J. Fried, M. D. Fox, K. Lee, P. Giacobbe, Z. J. Daskalakis, D. M. Blumberger, J. Downar, Concordance between BeamF3 and MRIneuronavigated target sites for repetitive transcranial magnetic stimulation of the left dorsolateral prefrontal cortex, Brain Stimulation 8 (5) (2015) 965–973. doi: 10.1016/j.brs.2015.05.008.

[4] K. A. Caulfield, H. H. Fleischmann, C. E. Cox, J. P. Wolf, M. S. George, L. M. McTeague, Neuronavigation maximizes accuracy and precision in TMS positioning: Evidence from 11,230 distance, angle, and electric field modeling measurements, Brain Stimulation 15 (5) (2022) 1192–1205. doi:10.1016/j.brs.2022.08.013.

[5] T. Herbsman, D. Avery, D. Ramsey, P. Holtzheimer, C. Wadjik, F. Hardaway, D. Haynor, M. S. George, Z. Nahas, More lateral and anterior prefrontal coil location is associated with better repetitive transcranial magnetic stimulation antidepressant response, Biological Psychiatry 66 (5) (2009) 509–515. doi:10.1016/j.biopsych.2009.04.034.

[6] A. T. Sack, R. Cohen Kadosh, T. Schuhmann, M. Moerel, V. Walsh, R. Goebel, Optimizing functional accuracy of TMS in cognitive studies: A comparison of methods, Journal of Cognitive Neuroscience 21 (2) (2009) 207–221. doi:10.1162/jocn.2009.21126.

[7] K. A. Johnson, M. Baig, D. Ramsey, S. H. Lisanby, D. Avery, W. M. McDonald, X. Li, E. R. Bernhardt, D. R. Haynor, P. E. Holtzheimer, H. A. Sackeim, M. S. George, Z. Nahas, Prefrontal rTMS for treating depression: Location and intensity results from the OPT-TMS multi-site clinical trial, Brain Stimulation 6 (2) (2013) 108–117. doi:10.1016/j.brs.2012.02.003.

[8] P. B. Fitzgerald, Targeting repetitive transcranial magnetic stimulation in depression: do we really know what we are stimulating and how best to do it?, Brain Stimulation 14 (3) (2021) 730–736. doi: 10.1016/j.brs.2021.04.018.

[9] L. Ning, N. Makris, J. A. Camprodon, Y. Rathi, Limits and reproducibility of resting-state functional MRI definition of DLPFC targets for neuromodulation, Brain Stimulation 12 (1) (2019) 129–138. doi:10.1016/j.brs.2018.10.004.

[10] V. A. Cardenas, J. V. Bhat, A. M. Horwege, T. J. Ehrlich, J. Lavacot, D. H. Mathalon, G. H. Glover, B. J. Roach, B. W. Badran, S. D. Forman, M. S. George, M. E. Thase, J. A. Yesavage, D. Yurgelun-Todd, A. C. Rosen, Anatomical and fMRI-network comparison of multiple DLPFC targeting strategies for repetitive transcranial magnetic stimulation treatment of depression, Brain Stimulation 15 (1) (2022) 63–72. doi:10.1016/j.brs.2021.11.008.

[11] C. Nauczyciel, P. Hellier, X. Morandi, S. Blestel, D. Drapier, J. C. Ferre, C. Barillot, B. Millet, Assessment of standard coil positioning in transcranial magnetic stimulation in depression, Psychiatry Research 186 (2-3) (2011) 232–238. doi:10.1016/j.psychres.2010.06.012.

[12] S. M. Goetz, T. Kammer, Neuronavigation, in: E. M. Wassermann, A. V. Peterchev, U. Ziemann, S. H. Lisanby, H. R. Siebner, V. Walsh (Eds.), The Oxford Handbook of Transcranial Stimulation, 2nd Edition, Oxford University Press, Oxford, 2023, pp. 183–226. doi:10.1093/oxfordhb/9780198832256.013.7.

[13] T. S. Riis, S. Lunt, J. Kubanek, MRI-free targeting of deep brain structures based on facial landmarks, Brain Stimulation 18 (1) (2025) 131–137. doi:10.1016/j.brs.2024.12.1478.

[14] A. E. Nieminen, J. O. Nieminen, M. Stenroos, P. Novikov, M. Nazarova, S. Vaalto, V. V. Nikulin, R. J. Ilmoniemi, Accuracy and precision of navigated transcranial magnetic stimulation, Journal of Neural Engineering 19 (6) (2022) 066037. doi:10.1088/1741-2552/aca71a.

[15] R. Elfring, M. de la Fuente, K. Radermacher, Assessment of optical localizer accuracy for computer aided surgery systems, Computer Aided Surgery 15 (1–3) (2010) 1–12. doi:10.3109/10929081003647239.

[16] N. Matilainen, J. Kataja, I. Laakso, Verification of neuronavigated TMS accuracy using structured-light 3D scans, Physics in Medicine and Biology 69 (8) (2024) 085004. doi:10.1088/1361-6560/ad33b8.

[17] J. Ruohonen, J. Karhu, Navigated transcranial magnetic stimulation, Neurophysiologie clinique/Clinical neuro-physiology 40 (1) (2010) 7–17.

[18] R. R. Shamir, L. Joskowicz, S. Spektor, Y. Shoshan, Localization and registration accuracy in image guided neurosurgery: a clinical study, International Journal of Computer Assisted Radiology and Surgery 4 (2009) 45–52. doi:10.1007/s11548-008-0268-8.

[19] A. I. Omara, M. Wang, Y. Fan, Z. Song, Anatomical landmarks for point-matching registration in image-guided neurosurgery, International Journal of Medical Robotics and Computer Assisted Surgery 10 (2014) 55–64. doi:10.1002/rcs.1509.

[20] J. M. Fitzpatrick, J. B. West, C. R. Maurer, Predicting error in rigid-body point-based registration, IEEE Transactions on Medical Imaging 17 (5) (1998) 694–702. doi:10.1109/42.736021.

[21] L. G. Farkas, Anthropometry of the Head and Face, 2nd Edition, Raven Press, New York, 1994.

